# Comparative evaluation of bioinformatic tools for virus-host prediction and their application to a highly diverse community in the Cuatro Ciénegas Basin, Mexico

**DOI:** 10.1101/2023.08.31.555565

**Authors:** Alejandro Miguel Cisneros-Martínez, Ulises E. Rodriguez-Cruz, Luis D. Alcaraz, Arturo Becerra, Luis E. Eguiarte, Valeria Souza

**Author notes:** Corresponding author (VS).

## Abstract

The sheer diversity of unculturable viruses has prompted the need to describe new viruses through culture-independent techniques. The associated host is one important phenotypic feature that can be inferred from metagenomic viral contigs -- thanks to the development of various bioinformatic tools. Here we compare the performance of recently developed tools for virus-host prediction on a dataset of 1,046 virus-host pairs and then apply the best-performing tools on a metagenomic dataset derived from a highly diverse transiently hypersaline site known as Archaean Domes within the Cuatro Ciénegas Basin, Coahuila, Mexico. We also introduce a virus-host prediction tool called CrisprCustomDB, which uses specific criteria to solve controversial host assignments with custom spacers databases. Host-dependent alignment-based methods showed an average precision of 83% and a sensitivity from 13.7% to 17.7%, whereas host-dependent alignment-free methods achieved an average precision of 75.7% and a sensitivity of 57.5%. RaFAH, a virus-dependent alignment-based tool, had the best performance overall (F1_score = 95.7%). However, when applied to the highly diverse metagenomic dataset, the host-dependent alignment-based (*e.g.*, CrisprCustomDB) and alignment-free (*e.g.*, PHP) methods showed the greatest agreement with each other, even though they are fundamentally different methods. This is because instead of depending on known hosts or viruses-with-known-host databases, they can directly relate metagenomic viral contigs and metagenome-assembled genomes from the same dataset. Such methods also showed the greatest consistency between the source environment and the predicted host taxonomy, habitat, lifestyle, or metabolism, revealing that Archaean Domes viruses likely infect halophilic Archaea as well as a variety of Bacteria which may be halophilic, halotolerant, alkaliphilic, thermophilic, oligotrophic, sulfate-reducing or marine-related. Consequently, using a combination of methods and qualitative validations relating to the source environment and the predicted host biology will increase the number of correct predictions, mainly when dealing with novel viruses.

## Introduction

Until 1990, the International Committee on Taxonomy of Viruses (ICTV) requested detailed information on biological properties to describe and classify new viruses. These biological properties were observed either *in vitro* (*e.g.*, replication cycle, virion structure and antigenic relationships) or in natural host interactions (*e.g.*, pathogenicity, epidemiology, and host range). However, with the development of DNA sequencing techniques, bioinformatics tools, and methods to study molecular evolution, the need to incorporate genomic information -- specifically phylogenetic groupings--to classify novel viruses has arisen [1]. In addition, the development of these tools now allows metagenomic analyses that can detect up to 90% of unknown viruses (non-culturable) in various environments. For example, the TARA ocean expedition assembled viruses from different oceans worldwide and could only assign a family-level classification to 10% of the assembled viral contigs [2]. As a result, the scientific community has proposed incorporating metagenomic data into the ICTV taxonomy by using phenotypic traits and phylogenetic information inferred from assembled sequences and genomes [1].

The need to characterize viruses assembled from metagenomes has prompted various bioinformatic methods for virus-host prediction [3]. A recent study suggests a five-category classification [4] (Fig 1): i) host-dependent alignment-based methods; ii) host-dependent alignment-free methods; iii) virus-dependent alignment-based methods; iv) virus-dependent alignment-free methods; and v) integrative methods. Host-dependent alignment-based methods include methods based on homology signals, such as searching for homology between viral and host proteins, tRNAs, viral genomes and CRISPR spacers, integrated prophages, and protein-protein interactions (PPI). These methods are helpful for detecting recent infections but have the disadvantage that not all viruses share genes with their hosts, which tends to make them precise, but with a low detection rate [3]. CrisprOpenDB is a recently released tool that uses biological criteria to standardize host predictions based on CRISPR spacers with increased sensitivity and precision thanks to its >11 million spacers database derived from >300,000 candidate hosts [5].

**Fig 1.**
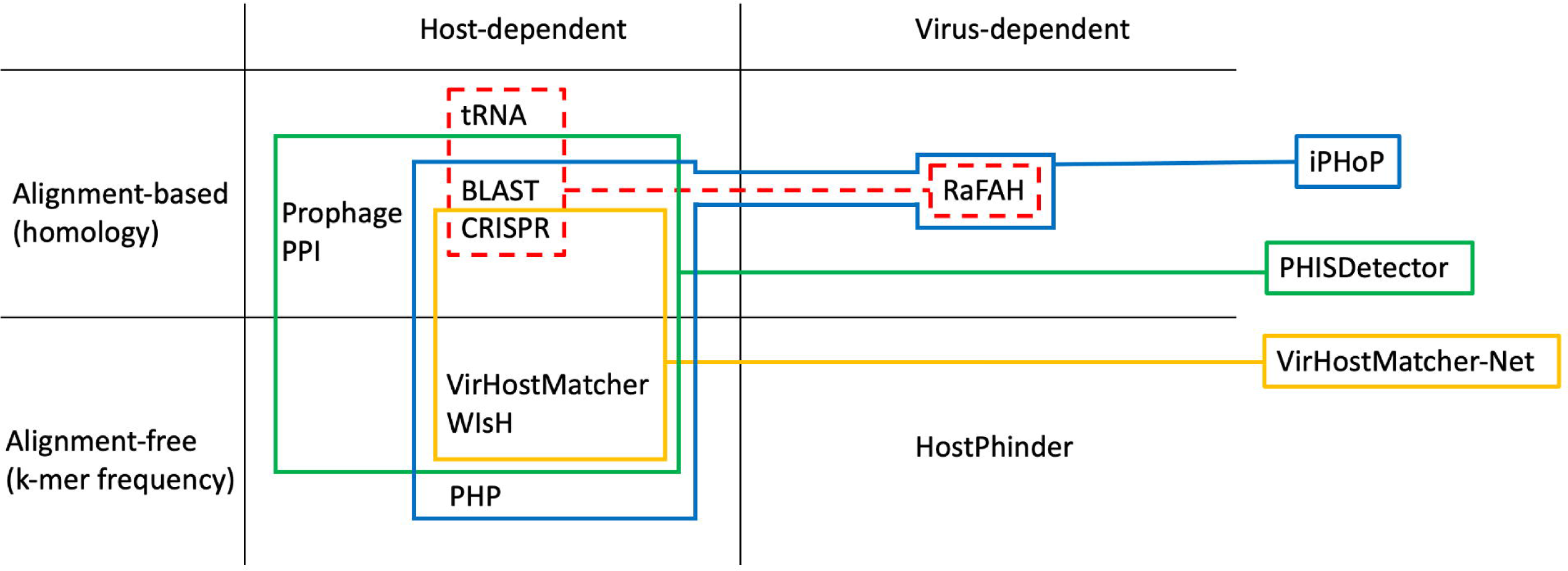
Classification of virus-host prediction methods. RaFAH uses host-dependent alignment-based methods to build part of its training database (red discontinuous lines). Integrative methods (iPHoP – blue lines; PHISDetector – green lines; VirHostMatcher-Net – yellow lines) attempt to exploit the virtues of a different number of methods. PPI = protein-protein interactions.

Host-dependent alignment-free methods include those based on sequence composition (*e.g.*, similarity in codon usage, similarity in oligonucleotide composition, and GC content), which rely on the notion that viruses, being genetic parasites, approximate their nucleotide composition to that of the host over time. This genomic mimicry may allow viruses to use the same tRNAs for protein synthesis or to evade the detection and degradation mechanisms of foreign nucleic acids. However, viruses can have similar sequence profiles independently, which can lead to a high false positive rate [3]. VirHostMatcher, which evaluates virus-host genome similarity through d*_2_ distance from 6-mer profiles [6], WIsH, which uses 8-mer profiles and Hidden Markov models (HMMs) [7] and PHP, which uses 4-mer profiles and a Gaussian model [8], are some well-known similar host-dependent alignment-free methods. These alignment-free strategies also include methods based on co-abundance profiles, which rely on the notion that viruses can only be found in the environment in which their host is also found. This profiling method requires the calculation of correlations of normalized abundance profiles of phage and bacteria in different environmental samples. However, they entail a major drawback: predator-prey interactions -- such as those described by the kill-the-winner model [9, 10] -- can generate positive or negative correlations, depending on where the interaction was at the time the sample was taken.

Instead of relying on host databases, virus-dependent methods depend on databases storing viruses with known hosts, to which query viruses are related either through homology signals (alignment-based) or their similarity in oligonucleotide composition (alignment-free). On the one hand, a machine-learning approach named Random Forest Assignment of Hosts (RaFAH) [11] is a virus-dependent alignment-based method that builds a part of its training database from CRISPR spacers, the presence of horizontally transferred genes and common tRNAs to ultimately associate the query virus to a virus with a known host through similarity in protein content. On the other hand, HostPhinder [12] is a virus-dependent alignment-free method that compares 16-mer profiles between query viruses and a database of 2,196 phages with known hosts.

Finally, integrative methods attempt to exploit the virtues of different methods like VirHostMatcher-Net [13], which integrates host-dependent alignment-based methods (CRISPR spacers) and host-dependent alignment-free methods (VirHostMatcher or WIsH) in a network framework, PHISDetector [14], which integrates BLAST [15], CRISPR spacers, prophage, and PPI analyses through a set of machine learning approaches, or iPHoP, which uses machine learning algorithms to compute taxonomy-aware scores for BLAST, CRISPR, VirHostMatcher, WIsH, and PHP, and integrates them with RaFAH results to obtain a final composite score [4].

Optimization of precision and sensitivity estimates within each approach has been achieved by either using more extensive reference databases (*e.g.,* CrisprOpenDB), by leveraging the power of different machine learning algorithms (PHP, RaFAH, VirHostMatcher-Net, PHISDetector, iPHoP), or by integrating different methods (VirHostMatcher-Net, PHISDetector, iPHoP). The publication of these tools is typically accompanied by validation tests with estimates of precision and sensitivity, as well as comparisons with other methods. However, most publications use different databases and sometimes use published values to compare the precision of different methods [6] directly. So far, Roux et al. [4] have compared the largest number of methods showing that host-dependent alignment-based methods can achieve high precision but suffer from low sensitivity. In contrast, host-dependent alignment-free methods have greater sensitivity but struggle to make correct predictions, while virus-dependent alignment-based methods such as RaFAH present both high sensitivity and precision. However, virus-dependent methods may underperform when predicting the host of novel viruses, which also affects, to a lesser extent, host-dependent alignment-based methods but not alignment-free methods [4]. Furthermore, we believe that the fact that host-dependent methods do not necessarily depend on a sizeable pre-compiled reference database (except for CrisprOpenDB) represents an advantage when predicting hosts of novel viruses since they allow to compare metagenomic viral contigs (mVCs) with potential hosts metagenome-assembled genomes (MAGs) from the same environment or even the same dataset.

Here, we present CrisprCustomDB, an algorithm inspired by CrisprOpenDB biological criteria to predict hosts based on CRISPR spacers, which allows the use of custom spacers databases (i.e., spacers predicted from the same metagenomic dataset as mVCs). Furthermore, we evaluated the precision and sensitivity of the different bioinformatic tools for virus-host prediction. Additionally, we applied some of the best performing tools on a metagenomic dataset derived from samples collected at a recently discovered pond known as Archaean Domes in the Cuatro Ciénegas Basin, Coahuila, in the north of Mexico [16, 17]. Archaean Domes is a seasonally fluctuating pond characterized by high pH and salinity, where an extreme diversity of Bacteria and Archaea has been recently described [16, 17]. In addition, its highly diverse viral community does not behave like those from other hypersaline or high pH sites in the face of environmental solid fluctuations [18], for which it is essential to characterize the virus-host relationships and interactions that may drive the microbial and viral diversity in such a unique site.

## Materials and Methods

### Benchmarking of bioinformatics tools for virus-host prediction

Genomes were selected by downloading three lists (S1_File): i) NCBI complete viral genomes (https://www.ncbi.nlm.nih.gov/genomes/GenomesGroup.cgi) filtered by host ‘bacteria’; ii) Virus-hostDB tabular report (https://www.genome.jp/ftp/db/virushostdb/) and; iii) RefSeq release 217 catalog (https://ftp.ncbi.nlm.nih.gov/refseq/release/release-catalog/), filtered by complete genomic molecule, not plasmid.

A link was established between the three tables so that if the viral genome accession had a match in the virus-hostDB, it was checked if the host taxa had a match in the RefSeq catalog (for a given virus with known host, check if the host has a complete genome). 1,029 phage genomes and 133 bacterial genomes were downloaded from NCBI (S2_File), which together made up 1,046 virus-host pairs with complete genomes. The performance of the virus-host prediction tools was evaluated at the genus level.

A custom script was written (https://github.com/AleCisMar/CrisprCustomDB/blob/main/benchmarking/compare_real-estimated.pl) to compare the estimated virus (accession)-host (genus) pairs with the actual virus (accession)-host (genus) pairs in order to obtain the true positives (TP), false positives (FP) and false negatives (FN) for every prediction tool. TP is the successful rejection of a null hypothesis denying any relationship between the real pairs. FP would typically be the incorrect rejection of a second null hypothesis about the relationship between falsely predicted or observed pairs (type I error). However, given that prediction tools are tested against a reference list of confirmed or expected virus-host pairs in this particular evaluation, FP is also incurring a type II error, which is the failure to reject that there is no relationship between the real pairs. Thus, FN includes FP and viruses with unassigned hosts (NA) (type II error).

Precision, sensitivity, and F1_score were calculated as follows:

Precision (Positive Predictive Value):

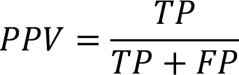

Sensitivity (True Positive Rate):

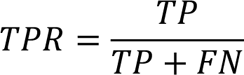

F1_score:

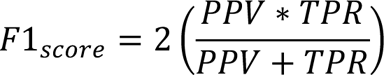

To run each program, we followed the instructions provided by the developers choosing the parameters for which they are reported to perform the best [5, 6, 7, 8, 11, 12, 13]. In addition, a Perl script inspired by CrisprOpenDB was developed (CrisprCustomDB available at https://github.com/AleCisMar/CrisprCustomDB). CrisprCustomDB uses the host assignment criteria of Dion et al. [5]. These are: (i) host if spacers have a maximum of 2 mismatches with the viral genome; (ii) if multiple candidates meet criteria 1, the one with the most spacers aligning with different regions of the viral genome is selected; (iii) if there are multiple candidates meeting criterion 2, the one with the spacer closest to the 5’ end in the array is selected; and (iv) if there are multiple candidates meeting criteria three the lowest shared taxonomic rank is assigned.

The CRISPR spacers were obtained with the CRISPRDetect tool [19] to test their performance on the custom database (133 bacterial genomes) with the parameters recommended by Dion et al. [5]. Since only 1,349 spacers (S2_File) were found in 40 of the 133 bacterial genomes (30%), two sensitivity calculations were made, one considering the 1,046 virus-host pairs and the other taking into account only the maximum number of virus-host pairs (261) that can be obtained given the number of possible hosts with spacers (40). CrisprCustomDB, VirHostMatcher, WIsH, and PHP were run on the custom database (1,029 phage genomes and 133 bacterial genomes).

Since HostPhinder, CrisprOpenDB, VirHostMatcher-Net, and RaFAH rely on large reference databases, only the 1,029 phage genomes were used as input. For PHP, an additional estimation was made using the reference database with 60,105 potential hosts provided by the authors. For VirHostMatcher, two estimations were made, both with a score ≤ 0.25. The first selects the most frequent host within the top 30 with the most similar profiles, and the second selects the most frequent host within the top 5 with the most similar profiles. For VirHostMatcher-Net, two estimations were also made. One without score restriction and the other limited to predictions with a score > 0.95.

### Virus-host prediction on assembled metagenomic reads Sample collection and sequencing

Sampling was carried out at the Archaean Domes of the Rancho Pozas Azules (26°49’41.9” N, 102°01’23.6” W), belonging to Pronatura Noreste, in the Cuatro Ciénegas Basin, Coahuila, in the North of Mexico, under SEMARNAT scientific permit number SGPA/DGVS/03121/15. Twelve samples were taken between 2016 and 2020. For microbial mats, seven surface samples (M1 – M6 and D0) were collected using a sterile scalpel dissection (8 cm^2^ / 40 cm^3^) and transferred to 50 mL conical tubes. 30 cm plastic tubes were used as sediment samplers to collect two additional microbial mat samples at 30 and 50 cm depth (D30 and D50). Three samples were collected at the shallow ellipsoid orange pools or orange circles (OC) [16, 17, 18]: one superficial water sample (C0) on a 50 mL conical tube and two more at depths of 30 and 50 cm (C30 and C50). All samples were stored in liquid nitrogen until processing.

DNA was extracted according to [20] at the Laboratorio de Evolución Molecular y Experimental of the Instituto de Ecología, Universidad Nacional Autónoma de México, in Mexico City. Briefly, the extractions followed a column-based protocol with a Fast DNA Spin Kit for Soil (MP Biomedical) [21]. Total DNA was sent to CINVESTAV-LANGEBIO, Irapuato, México, for shotgun sequencing with Illumina Mi-Seq paired-end 2×300 technology.

All sequence reads are available on the National Centre for Biotechnology Information (NCBI) Sequence Reads Archive (SRA) under the BioProject accession: PRJNA847603.

### Read processing and assembly of mVCs

The read quality was assessed with FastQC v0.11.9 [22]. Adapter removal and quality filtering were performed with Trimmomatic v0.39 [23] using a sliding window of 4 base pairs excluding reads with an average quality of less than 30 and less than 20 nucleotides. Clean reads were assembled with SPAdes 3.15.2 [24] using the --metaviral option. The viralVerify and viralComplete scripts (included in the SPAdes package) were used to verify that the assembled contigs correspond to viral genomes and to assess genome completeness, respectively. The circularity of the viral contigs was checked. When necessary, the position of sequences was adjusted prior to gene prediction and annotation with the help of custom scripts (available at https://github.com/AleCisMar/GenomicTools) that make use of BLAST [15], EMBOSS [25], Prodigal [26], and HMMER [27].

### Read processing, assembly, and taxonomic assignment of MAGs

The quality of the raw data was assessed with FastQC (v0.11.8) [22] and filtered with Trimmomatic (v0.39) [23]. The reads were then assembled using MetaSPAdes (v3.15.3) [28], and the contigs obtained in the assembly were used to perform read binning or clustering, which was performed with MaxBin2 (v2.2.7) [29] and MetaBat2 (v2.12.1) [30]. Binning refiner (v1.4.2) software [31] was used to reduce the percentage of contamination in the bins. The integrity of the MAGs was assessed using CheckM (v1.1.3) [32] with the default settings.

For taxonomic assignment and placement of MAGs on the phylogenetic tree of life, we used the program GTDB-tk (v1.6.0) [33], which identifies 122 and 120 marker genes of archaea and bacteria, respectively, using HMMER [27]. Briefly, genomes are assigned to the domain with the most identified marker genes. Selected domain-specific markers are aligned with HMMER, concatenated into a single multiple sequence alignment, and trimmed with the ∼5000-column Bacteria or Archaea mask used by GTDB [33].

### Implementation of virus-host prediction tools on metagenomic data

After taxonomic classification of the MAGs, prediction of the spacer sequences of the CRISPR arrays found in the MAGs was performed using the CRISPRCasTyper program (v 1.3.0) [34] using the following parameters cctyper -t 4 --prodigal single –circular. 2,660 spacers (S3_File) were found. The mVCs were run against the spacer database with Blastn [15], allowing for a maximum of 2 mismatches. Spacers were predicted with the CRISPRDetect tool [19] to implement CrisprCustomDB pipeline using array_quality_score_cutoff of 3, as recommended for FASTA files. This spacers prediction for CrisprCustomDB analysis resulted in 1,062 spacers (S3_File). The mVCs were also run with CrisprOpenDB, which uses an 11,674,395 spacers database [5]. Virus-host predictions were also made with RaFAH [11] and PHP [8]. For PHP, k-mer frequencies were calculated for all MAGs (S3_File).

## Results

### Benchmarking of bioinformatics tools for virus-host prediction

The best three performing tools for complete bacteria and phage genomes datasets (F1_score) were RaFAH, PHP, and VirHostMatcher-Net. They were followed by WIsH, VirHostMatcher, CrisprOpenDB, HostPhinder, and CrisprCustomDB at the bottom (Table 1).

**Table 1.**
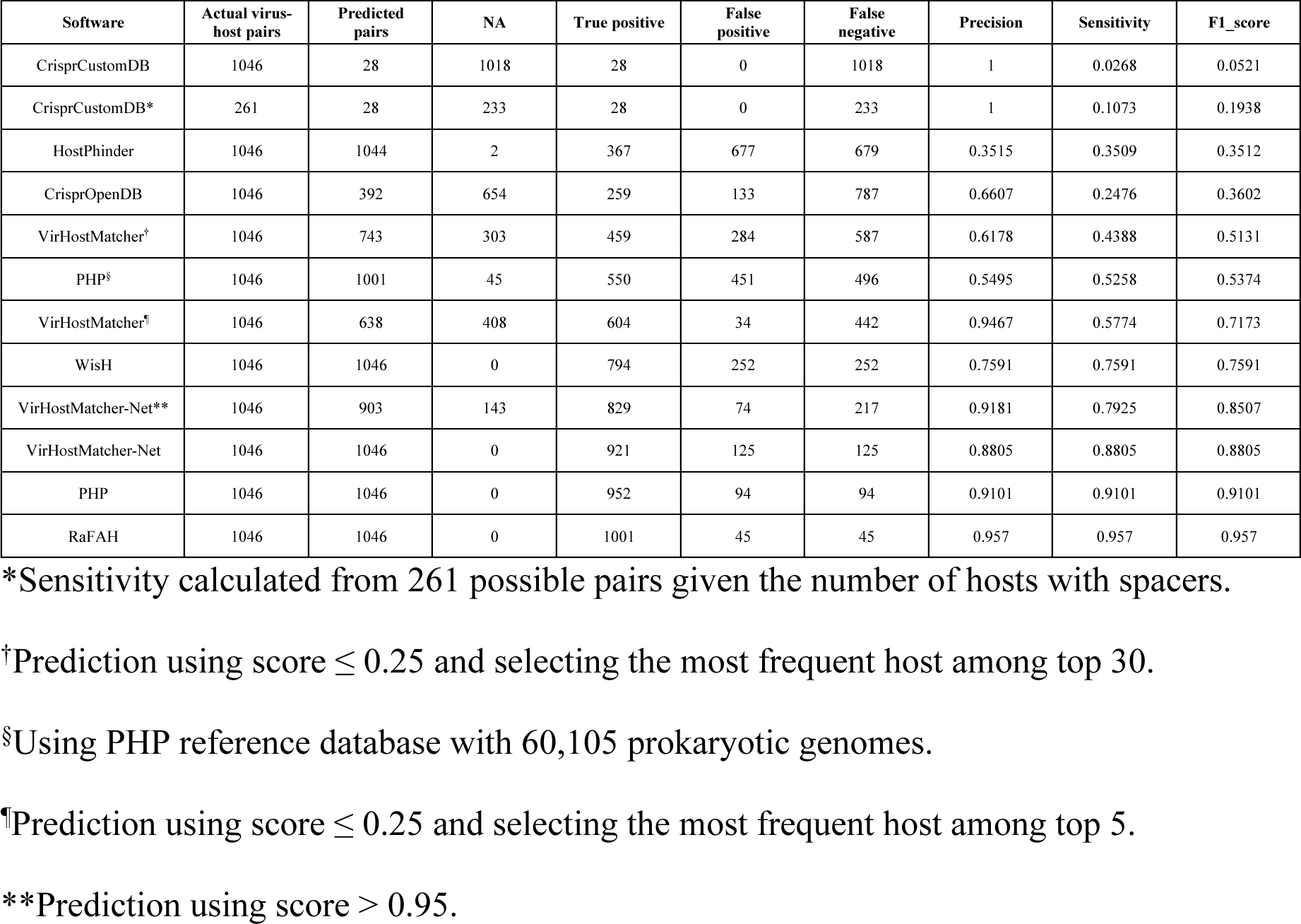
Precision, sensitivity, and F1_score estimates of the different virus-host prediction tools.

CrisprOpenDB made 392 predictions, of which 259 were correctly estimated. These results translate into a sensitivity of 24.76%, a precision of 66.07%, and an F1_score of 36.02% (Fig 2). On the other hand, CrisprCustomDB only predicted 28 pairs, all of which were correct (precision = 100%). Considering that we are benchmarking on 1,046 virus-host pairs, CrisprCustomDB reached a sensitivity of only 2.68% and an F1_score of 5.21%. It is important to note that spacers were identified in only 30% of the bacterial genomes in the custom database. Consequently, the maximum number of predicted pairs was 261. This consideration increases the sensitivity to 10.73% and the F1_score to 19.38% (Fig 2).

**Fig 2.**
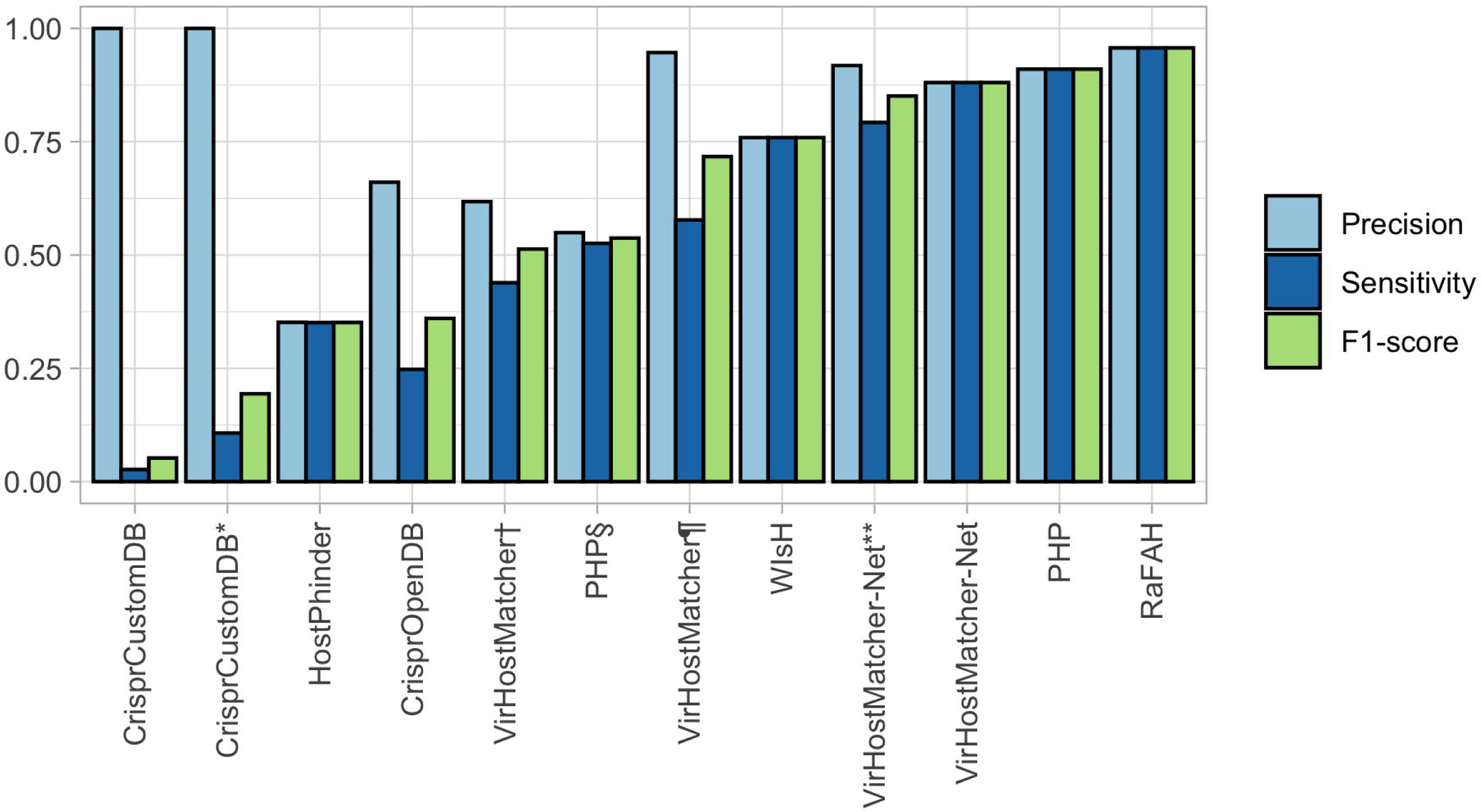
Precision, sensitivity, and F1_score estimates of the different virus-host prediction tools. For CrisprCustomDB*, sensitivity was estimated considering 261 possible pairs. VirHostMatcher† was tested with a score ≤ 0.25 and selected the most frequent host within the top 30. VirHostMatcher¶ was tested with the same parameters but selecting the most frequent host within the top 5. PHP§ was tested against a reference database of 60,105 potential hosts. For VirHostMatcher-Net**, only predictions with a score > 0.95 were kept.

Alignment-free methods evaluated here make predictions by comparing the oligonucleotide profile of a virus to either the oligonucleotide profile of viruses with a known host (HostPhinder) or the oligonucleotide profile of bacteria (VirHostMatcher, WIsH, PHP). Although HostPhinder predicted 1,044 pairs, most predictions were incorrect (677). Hence, it had the lowest performance of the alignment-free methods (Fig 2), with a sensitivity of 35.09%, a precision of 35.15%, and an F1_score of 35.12%.

VirHostMatcher was executed with two different criteria: i) selecting the most frequent host among the top thirty and; ii) selecting the most frequent host among the top five. When using the first criterion, VirHostMatcher generated more predictions (743 compared to 638) and produced more false positives (284 compared to 34). As a result, it achieved lower sensitivity (43.88% compared to 57.74%), precision (61.78% compared to 94.67%), and F1_score (51.31% compared to 71.73%).

Among these methods, WIsH and PHP emerged as the top predictors, achieving the maximum number of pairs (1,046). WIsH demonstrated a sensitivity, precision, and F1_score of 75.91%, whereas PHP appeared as the best-performing alignment-free method (Fig 2) with a sensitivity, precision, and F1_score of 91.01%. PHP was also tested against a reference database with 60,105 potential hosts provided by the authors. However, this test resulted in fewer predictions (1,001) and lower sensitivity (52.58%), precision (54.95%), and F1_score (53.74%).

VirHostMatcher-Net was executed using two approaches: first, by setting a prediction threshold with a score > 0.95 and, second, without any score restrictions. Restricting the final host assignment to predictions with higher scores resulted in higher accuracy (91.81% vs. 88.05%) at the expense of lower sensitivity (79.25% vs. 88.05%) and, as a consequence, a lower F1 score (85.07% vs. 88.05%). Meanwhile, RaFAH achieved an accuracy, sensitivity, and F1 score of 95.70%, making it the algorithm with the best overall performance (Fig 2).

### Virus-host predictions on assembled metagenomic reads from Archaean Domes, Cuatro Ciénegas Basin, Mexico

To predict the host of mVCs from Archaean Domes at Cuatro Ciénegas Basin, based on CRISPR spacers predicted on MAGs from the same dataset, we employed two related approaches. The first approach involved conducting a Blastn search using 2,660 spacers, with a maximum of 2 mismatches as the only criterion. The second approach involved CrisprCustomDB, using 1,062 spacers to solve problematic host assignments. Additionally, we performed predictions using CrisprOpenDB, PHP, and RaFAH, as these tools demonstrated superior performance in their respective categories. Since HostPhinder showed lower performance than all other alignment-free methods, and PHP and RaFAH outperformed VirHostMatcher-Net, we did not make predictions with these tools. While PHP was executed on the Archaean Domes MAGs, CrisprOpenDB and RaFAH only required the mVCs, as they relied on their reference databases.

The ordinary CRISPR approach resulted in eight predictions (Table 2). Half of the mVCs (C50N1L42, C0N5L506, M1N5L607, and C9N1L394) were assigned to hosts in the phylum *Desulfobacterota*. The other half were associated with bacteria of the class *Gammaproteobacteria.* Two mVCs (M5N2L438 and M6N1L439) were predicted to infect bacteria of the genus *Halorhodospira*. One contig, the M4NL642, was assigned to the genus *Halochromatium,* and the contig C30N1L64, was assigned to two possible hosts: *Thiohalorhabdus* or *Thiohalospira*.

**Table 2.**
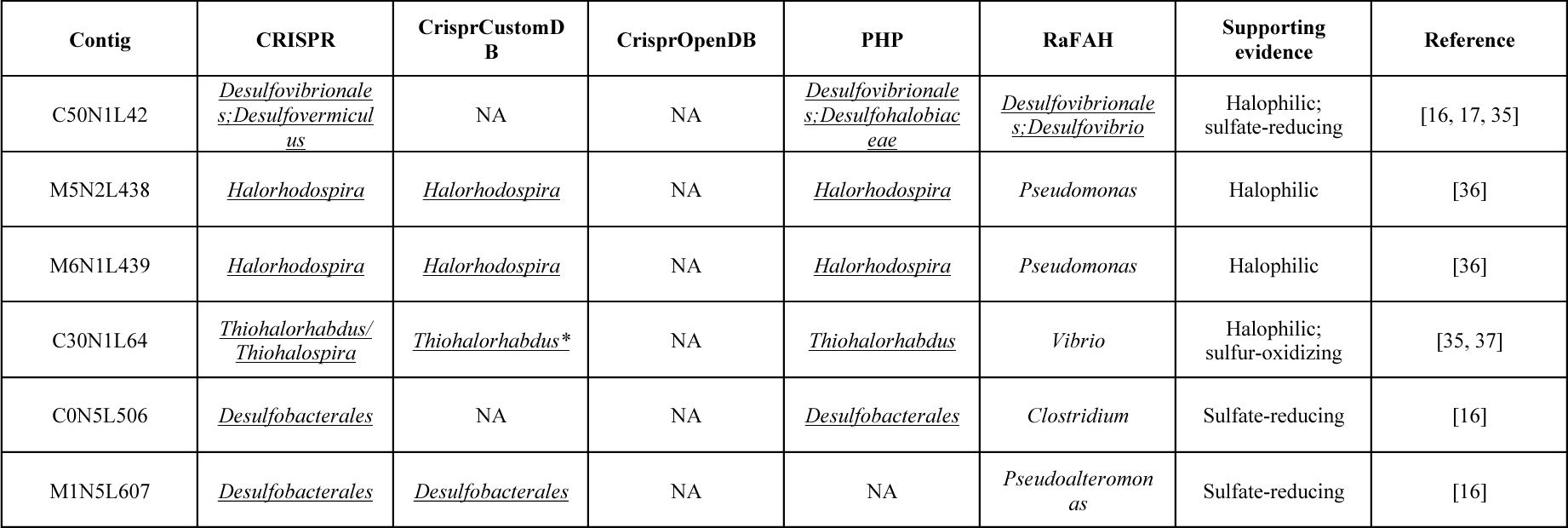
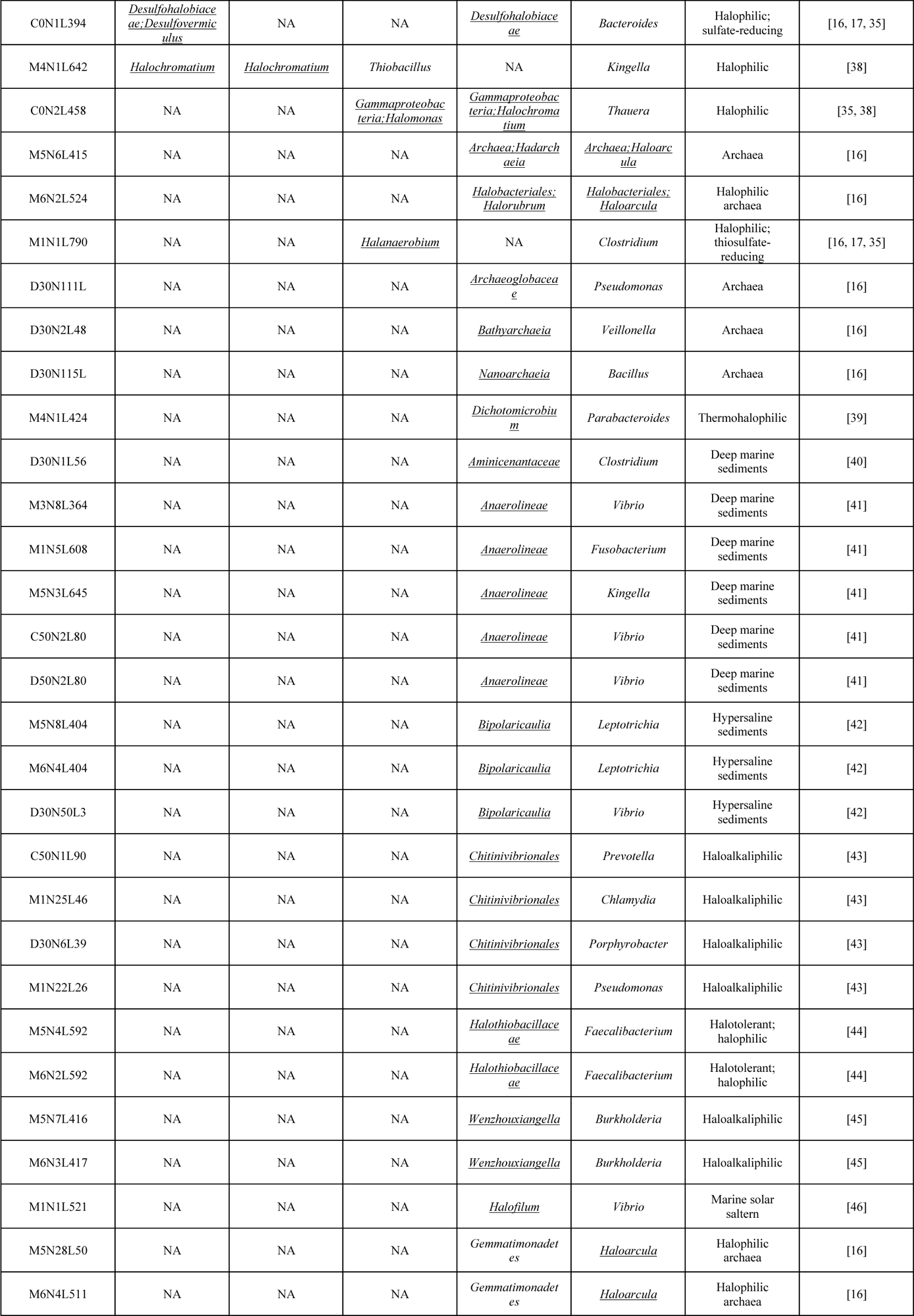
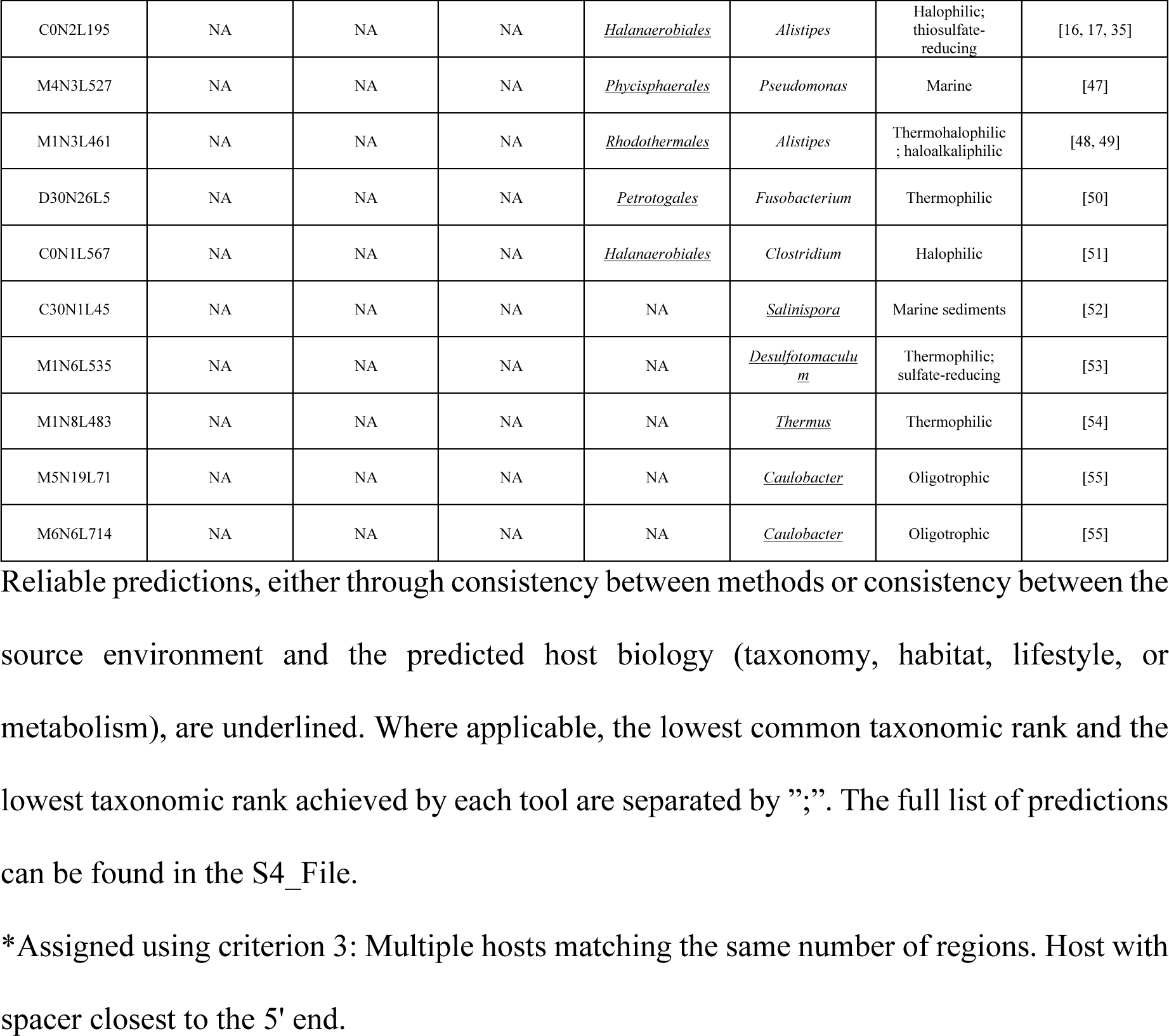
46 host predictions on mVCs from Archaean Domes Pond, Cuatro Ciénegas, Mexico, designated as reliable according to different criteria.

CrisprCustomDB made five predictions, all consistent with those made by the ordinary CRISPR approach. These included one of the *Desulfobacterota* and three *Proteobacteria*. The fifth prediction assigned contig C30N1L64 to *Thiohalorhabdus* for being the host with the spacer closest to the 5’ end, solving the problem of two possible hosts (Table 2). CrisprOpenDB made five predictions (S4_File). The only contig for which all three CRISPR-based methods made a prediction is contig M4N1L642. However, ordinary CRISPR and CrisprCustomDB predicted it to infect the proteobacteria *Halochromatium*, while CrisprOpenDB predicted *Thiobacillus* as the phage host. CrisprOpenDB predicted contig C0N2L458 to infect *Gammaproteobacteria* of the genus *Halomonas* (Table 2).

PHP made 54 predictions. Three of the contigs assigned to *Desulfobacterota* by the ordinary CRISPR approach (C50N1L42, C0N5L506, and C9N1L394) were equally assigned by PHP. Also, three contigs assigned to *Proteobacteria* in both the ordinary CRISPR and CrisprCustomDB approaches were independently assigned the same by PHP. This concordance includes contigs M5N2L438 and M6N1L439, assigned to *Halorhodospira*, and contig C30N1L64, which was also assigned to *Thiohalorhabdus*. Additionally, PHP agreed with CrisprOpenDB on the host assignment for contig C0N2L458 at the class level but suggestd it infects bacteria of the genus *Halochromatium* instead of *Halomonas* (Table 2).

RaFAH produced 87 predictions, of which only three were supported by the other methods. These included the host assignment for contig C50N1L42 (*Desulfobacterota*), which is consistent with the ordinary CRISPR approach and PHP, and the host assignment for contig M5N6L415, which is consistent with PHP in predicting it to infect Archaea. The most similar assignment was observed for contig M6N2L524, predicted to infect *Euryarchaeota* of the genus *Haloarcula* by RaFAH and *Euryarchaeota* of the genus *Halorubrum* by PHP (Table 2). Finally, RaFAH was the only method that correctly predicted the host of *Escherichia* virus ΦX174, which was used as a positive control for DNA sequencing (S4_File).

## Discussion

The increasing number of virus-host prediction tools prompted us to perform a comparative evaluation of the most popular and recently released tools (Fig 1). Unfortunately, due to computational limitations related to the size of the databases, we could not evaluate either PHISDetector [14] or iPHoP [4]. Since there is a discrepancy in the performance of PHISDetector compared to VirHostMatcher-Net [4, 14], we can only conclude which tool performs the best once we compare them under the same methodological framework. As for iPHoP, this is probably the best-performing integrative tool [4], as it integrates RaFAH into its host prediction algorithm, which has shown better performance than VirHostMatcher-Net both here (Fig 2) and in its original publication [11].

According to the literature, it is understood that following iPHoP, PHISDetector (compared with VirHostMatcher-Net, PHP, WIsH and VirHostMatcher) [14] and RaFAH (reported with higher F1 score than the combination of CRISPR, BLAST and tRNAs, followed by VirHostMatcher-Net, WIsH, HostPhinder, and CRISPR, BLAST and tRNAs individually) [11] are the most precise tools. They are likely to be followed by VirHostMatcher-Net (more precise than similarity networks, CRISPR, BLAST, WIsH, and VirHostMatcher) [13], PHP (reported less precise than CRISPR and BLAST, which, however, have very low sensitivity, but are more precise than WIsH and VirHostMatcher) [8] and WIsH (reported to be more precise than VirHostMatcher, especially for incomplete or short viral genomes) [7]. Lastly, CrisprOpenDB (reported to have similar precision to WIsH) [5], HostPhinder (reported to be more precise than BLAST) [12], and VirHostMatcher (compared to values published by Edwards et al. [3] appears to have similar precision to homology methods (BLAST, prophage, and CRISPR) and higher than early implementations of the k-mer method, abundance profiling and GC content) [6] appear to be the least precise tools.

To test the above interpretations about the performance of virus-host prediction tools, we downloaded 1,029 and 133 complete phages and bacterial genomes, respectively. (S1_File and S2_File), making up 1,046 virus-host pairs. We did not use Archaea viruses and their respective hosts, because we could only retrieve eight pairs following the method described in the Materials and Methods section. In addition, some of the virus-host prediction tools evaluated here are explicitly trained on Bacteria and their corresponding phages (*e.g.* [5]) and, therefore, cannot be used to evaluate thier performance on viruses of Archaea. The performance of the virus-host prediction tools was evaluated at the genus level because performance comparisons are often consistent across taxonomic ranks [6, 7, 8, 11, 13, 14], and because it may be more biologically informative than higher taxonomic rank predictions.

As noted elsewhere [3, 4, 8], CRISPR-based methods demonstrated high precision at the expense of sensitivity. In our study, CrisprCustomDB achieved 100% precision but only made 28 out of 1,046 predictions, resulting in extremely low sensitivity. This limitation may be attributed to only 30% of the bacterial genomes having spacers, automatically excluding the remaining 70% from host predictions. Even when considering the maximum number of virus-host pairs that can be predicted given the number of hosts with spacers, sensitivity remained the lowest of all compared methods. Alternatively, the ability of CRISPR-based methods to detect virus-host pairs may be hampered by the host sequence selection process (see Materials and Methods), which excludes plasmid sequences and, therefore, any CRISPR-Cas system possibly encoded therein [56].

As expected, the limited sensitivity of the CRISPR-based methods was overcome by CrisprOpenDB due to the use of a > 11 million spacers database, which made 364 more predictions while retaining a precision higher than some sequence compositions methods, making it a reliable alternative when only viral contigs are available. Although CrisprOpenDB has increased the sensitivity of CRISPR-based methods, they still need to catch up to newer sequence composition methods, which exhibit high sensitivity and improved precision. Such was the case of VirHostMatcher, WIsH, and PHP, which achieved sensitivity and precision > 50%. In contrast, HostPhinder had a higher sensitivity than CRISPR-based methods but the lowest precision among all compared methods. This result suggests that relying solely on transferring the host of the most similar virus may be a greedy and unreliable approach, especially when dealing with a highly diverse viral community with many unknown viruses.

VirHostMatcher did not perform better when assigning the most frequent taxon among a more significant number of possible hosts (up to 30) with a score ≤ 0.25, contrary to what has been reported [6]. Instead, using this consensus criterion among the top 30 scoring hosts yielded a precision even lower than that of CrisprOpenDB and WIsH, which is known to perform better with incomplete contigs [7], while assigning host among the top 5 reached the second highest precision overall. Such discrepancies may depend on the distribution of taxa within the studied dataset. For instance, while increasing the n possible hosts criterion, one can expect a higher probability of finding multiple high-scoring instances of a particular host only by chance on a highly diverse dataset.

PHP allows predictions to be made with custom databases and provides a database of 60,105 bacterial genomes within the program’s repository. Using this reference database, PHP obtains the second-lowest precision overall, while the custom database (133 bacterial genomes) elevated PHP as the most accurate and sensitive sequence composition method. This result implies that using an extensive reference database does not necessarily enhance the performance of virus-host prediction tools, unless the actual hosts are present. Thus, PHP may be a suitable tool, especially when working with MAGs and mVCs from the same metagenome. Also, although not directly tested, host-dependent alignment-free methods such as PHP were noticeably more effortless to set up and faster to execute than integrative methods and virus and host-dependent alignment-based methods.

RaFAH achieved the highest precision, sensitivity, and F1_score on the test data collection. However, only a couple of its predictions on the metagenomic dataset were consistent with those of CRISPR-based methods, PHP, or the environment from which the metagenomes were generated. The metagenomic data analyzed here came from samples taken within the Cuatro Ciénegas Basin which, despite being a desert oasis with oligotrophic waters, is known for sheltering diverse groups of microorganisms, many of which are endemic and related to marine microorganisms [57, 58]. Such diversity is believed to have evolved as a result of the long-standing environmental stability of a deep aquifer that recreates an ancient ocean conditions, and which nourishes the aquatic systems of Cuatro Ciénegas Basin through the movement of groundwater produced by the magmatic pouch deep in the Sierra San Marcos y Pinos [59]. Specifically, the environment from which samples were extracted is a shallow pond characterized by high pH and salinity known as Archaen Domes [16, 17, 18]. It has been shown that Archaean Domes harbors a great diversity of bacteria on a short spatial scale [17] and is one of the most diverse archaeal communities in the world [16]. Such diversity includes sulfate-reducing *Proteobacteria* and extreme halophilic *Euryarchaeota* [35]. In addition, a highly diverse viral community has recently been described where haloarchaeaviruses constitute an essential part [18]. Therefore, predictions pointing to halophilic Archaea, as well as halophilic, halotolerant, alkaliphilic, thermophilic, oligotrophic, sulfate-reducing, sulfur-oxidizing or marine Bacteria, were considered consistent with the environment in question (Table 2).

Although CrisprCustomDB was able to discriminate between possible hosts for contig C30N1L64 (further supported by PHP), the fact that the ordinary CRISPR approach made more predictions on the metagenomic dataset than CrisprCustomDB is likely reflecting the benefit of using a more extensive spacer database (see Materials and Methods) as previously discussed regarding the performance of CrisprOpenDB. However, the lack of consistency of CrisprOpenDB and RaFAH with the other methods suggests that relying on a >11 million spacers database [5] or on a Random Forest classifier based on the protein content of viruses with known host [11], respectively, may be beneficial only when the hosts or the assembled viruses are already known, or are closely related to hosts or viruses represented in the corresponding databases. Therefore, for highly diverse datasets likely to have a high proportion of novel viruses as the one tested here [18], it may be more appropriate to use host-based tools, either alignment-based or alignment-free, such as CrisprCustomDB or PHP, with *ad hoc* databases built with archaea and bacteria MAGs from the same dataset whenever possible.

Predictions on the metagenomic dataset show that fundamentally different methods such as CrisprCustomDB and PHP, can complement and support each other. Incorporating these tools along with RaFAH, the best-performing tool on the test dataset, in an integrative software such as iPHoP [4], allows tackling the host prediction problem from different angles, increasing the chance of making the correct predictions. Also, judging the predictions based on the consistency between the predicted host biology (*i.e.*, taxonomy, habitat, lifestyle, or metabolism) and the source environment of the query virus (Table 2) may provide additional validation, mainly when predicting hosts of novel viruses. However, some caution still needs to be exercised with this validation approach. For example, for predictions with less consistency between methods and at higher taxonomic ranks, there is an increased risk that the consistency between the source environment of mVCs and the biology of the predicted hosts will be rather ambiguous or even false.

Host prediction is one of the most critical features for characterizing mVCs, probably along with phylogenetic relationships. We wanted to know who the host is to learn more about the biology of the newly assembled virus, such as where it gets the resources to complete its replication cycle, what organisms it interacts with, and with whom it might co-evolve. However, although the host predictions presented here take us a step forward in characterizing Archaean Domes viruses, we still need to know the phylogenetic context, the evolutionary processes, and the functional adaptations that will allow us to better understand the origin of diversity at this particular site.

## Conclusions

The results presented here indicated that RaFAH, a virus-dependent alignment-based method, and PHP, a host-dependent alignment-free method, are the best-performing tools for virus-host prediction. Other methods showed different performances depending on the host selection criteria, scoring thresholds, and the reference database. It seems that CRISPR-based methods seem to benefit from using a more extensive spacers database when predicting hosts of already-known viruses. However, using a more extensive candidate host database did not enhance the performance of host-dependent alignment-free methods such as PHP.

The complementarity and support shown by CrisprCustomDB and PHP when executed on mVCs and MAGs from the same dataset, suggest that using such a combination of tools along with RaFAH may produce more reliable host assignments on highly diverse metagenomic datasets provided that predictions are consistent across multiple methods and the predicted host taxonomy, habitat, lifestyle, or metabolism is consistent with the source environment.

Finally, host predictions on mVCs from Archaean Domes showed that viruses inhabiting such environment infect halophilic Archaea as well as a variety of Bacteria which may be halophilic, halotolerant, alkaliphilic, thermophilic, oligotrophic, sulfate-reducing or marine-related. These predictions are consistent with the particular environment and the known geological and biological evolution of the Cuatro Ciénegas Basin and its microorganisms.

## Acknowledgments

We greatly acknowledge Dr. Erika Aguirre-Planter, Dr. Rosalinda Tapia, Dr. Laura Espinosa-Asuar, Jazmín Sánchez-Pérez, Mariette Viladomat-Jasso, Rodrigo Vázquez, and Manuel Rosas from the Instituto de Ecología, UNAM, for their logistical and technical support and assistance during sample collection, and laboratory processing. We also acknowledge Dr. José Alberto Campillo-Balderas for reviewing the manuscript, and Rodrigo García-Herrera for providing access to the LANCIS-Instituto de Ecología’s Supercomputer Unit at UNAM. Finally, we thank SEMARNAT and APFF Cuatro Ciénegas for facilitating the sampling and, in particular, Rancho Pozas Azules, PRONATURA Noreste for access and permission to sample in the Cuatro Ciénegas Basin Natural Protected Area.

## Supporting information

**S1 File. Lists of NCBI complete bacterial virus genomes and RefSeq complete bacterial genomes used for benchmarking virus-host prediction tools.** The file contains six sheets—the first one lists bacteriophage genomes. The second is the Virus-Host DB table. The third one is the RefSeq release catalog with complete bacterial genomes—the fourth lists the virus-host pairs, including their accessions, used for benchmarking. The fifth sheet is the reference accession-genus list against which each prediction was compared. The sixth and final sheet contains the prediction results of each tool, using the parameters on which they perform the best.

**S2 File. NCBI complete bacterial virus genomes and RefSeq complete bacterial genomes sequence files used for benchmarking virus-host prediction tools.** It includes viral and bacterial genomes in fasta format. It also contains files with predicted CRISPR arrays and spacers.

**S3 File. Files used for testing virus-host prediction tools on metagenomic data from Archaean Domes, Cuatro Ciénegas Basin, Mexico.** It includes spacers needed to run predictions with BLAST and CrisprCustomDB and the host k-mer file needed to run predictions with PHP.

**S4 File. Host prediction results on mVCs from Archaean Domes, Cuatro Ciénegas Basin, Mexico.** It includes a table with contig information and host predictions and a Venn diagram showing the number of shared predictions by the different tools. Contigs highlighted with beige backgrounds will likely represent the same virus according to their protein domain content and host prediction. A red line delimits unreliable predictions from predictions supported by only one method but with consistency between the predicted host biology and the virus source environment. Above the yellow line, predictions are supported by two methods. Above the green line, predictions are supported by three methods.

## Notes

### Competing Interest Statement

I have read the journal's policy and the authors of this manuscript have the following competing interests: LDA is an Academic Editor for this journal. This does not alter our adherence to the PLOS ONE policy on sharing data and materials.

